# Fin whale singalong: evidence of song conformity

**DOI:** 10.1101/2022.10.05.510968

**Authors:** Miriam Romagosa, Sharon Nieukirk, Irma Cascão, Tiago A. Marques, Robert Dziak, Jean-Yves Royer, Joanne O’Brien, David K. Mellinger, Andreia Pereira, Arantza Ugalde, Elena Papale, Sofia Aniceto, Giuseppa Buscaino, Marianne Rasmussen, Luis Matias, Rui Prieto, Mónica A. Silva

**Affiliations:** Okeanos - Instituto de Investigação em Ciências do Mar, Universidade dos Açores & IMAR – Instituto do Mar, Horta, Portugal; Cooperative Institute for Marine Ecosystem and Resources Studies, Oregon State University, Oregon, USA; Centre for Research into Ecological and Environmental Modelling, University of St Andrews, St Andrews, UK; Centro de Estatística e Aplicações, Departamento de Biologia, Faculdade de Ciências, Universidade de Lisboa, Lisboa, Portugal; NOAA Pacific Marine Environmental Laboratory, Hatfield Marine Science Center, Oregon, USA; Univ Brest, CNRS, Laboratoire Geosciences Ocean, Plouzane, France; Marine and Freshwater Research Centre, Galway-Mayo Institute of Technology, Galway, Ireland; Instituto Dom Luiz (IDL), Universidade de Lisboa, Lisboa, Portugal; Institute of Marine Sciences, ICM-CSIC, Barcelona, Spain; Institute for the Study of Anthropic Impacts and Sustainability in the Marine Environment of the National Research Council of Italy (CNR-IAS), Torretta Granitola, Italy; Akvaplan Niva, Tromsø, Norway; University of Iceland’s research center in Húsavík, Iceland

**Keywords:** Vocal learning, conformity, song evolution, fin whale, North Atlantic

## Abstract

Mechanisms driving song learning and conformity are still poorly known yet fundamental to understand the behavioural ecology of animals. Broadening the taxonomic range of these studies and interpreting song variation under the scope of cultural evolution will increase our knowledge on vocal learning strategies. Here, we analysed changes in fin whale (*Balaenoptera physalus*) songs recorded over two decades across the Central and Northeast Atlantic Ocean. We found a rapid (over 4 years) replacement of fin whale song types (different inter-note intervals - INIs) that co-existed with hybrid songs during the transition period and showed a clear geographic pattern. We also revealed gradual changes in INIs and note frequencies over more than a decade with all males adopting both rapid and gradual changes. These results provide evidence of vocal learning of rhythm in fin whale songs and conformity in both song rhythm and note frequencies.

## Introduction

Animal songs, used as acoustic sexual displays, can change and evolve as the result of cultural processes like vocal learning and conformity. These mechanisms play important roles in evolutionary diversification, population structure and demography^1,2^ and may even act as a ‘second inheritance system’ to genetic evolution^3^. Vocal learning is a form of social learning where animals modify their acoustic signals as a result of hearing and imitating conspecifics^4,5^. Janik and Slater (2000)^4^ distinguish vocal production learning, the ability to modify signals after experience with the signals of others, from usage learning, where an existing signal is used in a different context or sequence. When individuals are more likely to share song variants with nearby individuals than with more distant ones, we talk about conformity^6^.

The earliest evidence for vocal production learning and conformity in animals comes from male birds learning songs^7^ but a growing literature indicates that vocal learning is widespread across various singing species^8^. One example is the humpback whale (*Megaptera novaengliae*), whose complex songs differ across ocean regions^9^ and all males in a given area adopt song changes ^10,11^. These songs can undergo revolutions, where a population song type is rapidly replaced by a novel song type introduced from a neighbouring population^12^. Most authors agree that these spatial and temporal patterns in song changes can only be explained by vocal production learning^5,8,12,13^. Although similarities in song evolution patterns between taxa have led to fruitful comparative studies^2^, we are still far from understanding the functional mechanisms of vocal learning, how natural selection has shaped learning strategies ^14^, which are the benefits of conformity, or how individual decisions affect the dynamics of song evolution over larger temporal and social scales^14^. Additionally, most research to date has focused on complex songs and vocal learning of rhythm is still poorly understood, despite many species’ relying on rhythmic aspects of conspecifics’ signals^15,16^. A broader view including different mechanisms of vocal learning, from simple to complex, is needed to better understand the prevalence and evolution of these complex phenomena^5^.

The simplicity of fin whale (*Balaenoptera physalus*) songs^17^ compared to those of humpback whales or songbirds, offers an easier-to-interpret scenario that can provide new insights into the field of song cultural evolution. Fin whales produce low-frequency songs that are believed to act as mating displays^17^, because they are produced by males^18^ and intensify during the breeding season^17,19–21^. In this species, the song inter-note interval (INI) (i.e., rhythm) is the most distinctive parameter between regions^17,22–25^ and can change abruptly from one year to the next^22,24–26^ or progressively over time^26–29^. Moreover, frequencies of two fin whale song components, the 20-Hz note and the higher frequency (~130-Hz) upsweep (hereafter HF note)^22^, have been decreasing gradually over the last decade in different ocean basins^27,28^. So far, studies on fin whale song changes have been merely descriptive and only focused on the potential drivers of these variations^24–30^.

Here, we report the dynamics of temporal and spatial variation of fin whale song parameters (INIs and peak frequencies of two note types) in a wide area of the North Atlantic under the scope of cultural song evolution, which provide a unique opportunity to investigate unknown mechanisms of vocal behaviour in this species. Our work provides evidence of vocal learning of rhythm in fin whale songs, and conformity in both song rhythm and note frequencies. This is supported by our results showing: i) a rapid shift in song INIs across a vast area of the Northeast Atlantic in just four singing seasons with the existence of hybrid songs (including both INIs) during the transition period; ii) a clear geographic gradient of song types; iii) a gradual increase in INIs and a decrease in frequencies of two song components over more than a decade; and iv) the adoption of rapid and gradual changes by all males in a wide region. We finish up by discussing conformity in terms of song function and potential benefits for fin whales.

Investigating vocal learning and conformity in fin whale songs is essential to understand important aspects of fin whale behavioural ecology, such as how this species responds to changes in its acoustic environment, and how culture shapes these responses.

## Results

### Rapid song changes

Results showed a rapid change in INIs, across a vast area of the Oceanic Northeast Atlantic (ONA) region (Fig. 1A), where the 19s-INI song was completely replaced by the 12s-INI song in just four singing seasons (2000/2001 – 2004/2005) (Fig. 2A and B). By 2004, the 19s-INI song had disappeared from the SE ONA location (Fig. 2B) and was no longer detected in any of the sampled regions from 2007 to 2020, except from an isolated account in 2008 (Fig. 3A). During the transition period, both song types co-existed SE of the ONA region, with a notable percentage of hybrid songs containing both INIs (~24% hybrids in 2002/2003), and there was a clear spatial gradient across the entire ONA region in the prevalence of songs with each INI type (Figs. 2B and C). In 2002/2003, the 19s-INI song largely dominated in the SW ONA, with only 8% of hybrid songs. The proportion of 19s-INI songs decreased progressively to the east, reaching 0-8% in the easternmost locations (CE and NE), while the 12s-INI songs became prominent. Hybrid songs were more abundant (16-24%) at central ONA (NW, CW and SE) than in easternmost locations (NE and CE; ~12%) (Fig. 2C).

**Figure 1.**
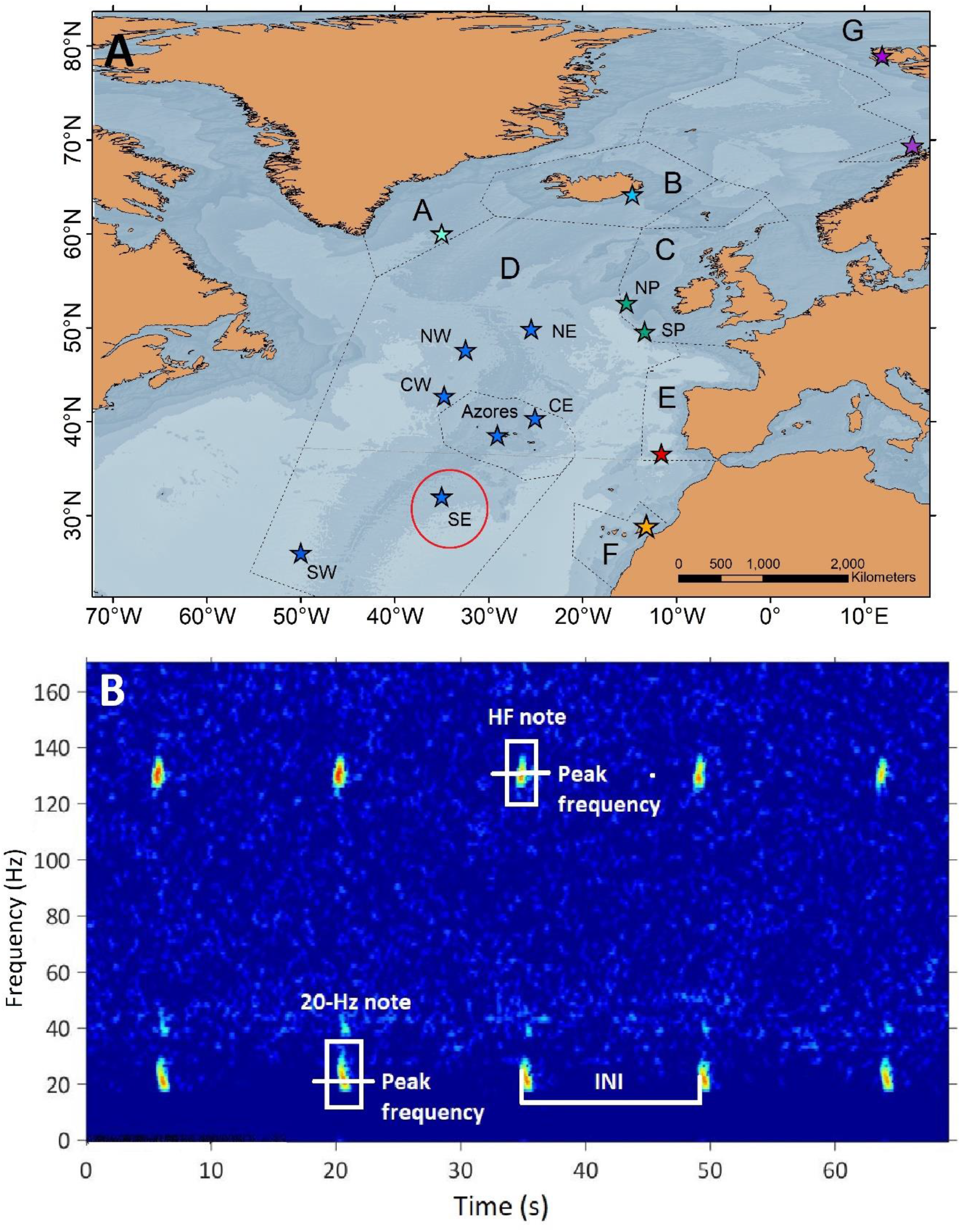
(A) Regions (dashed lines) and locations (stars) of acoustic recordings. Regions were delimited according to ICES ecoregions. A: Greenland Sea, B: Icelandic Waters, C: Celtic Sea, D: Oceanic Northeast Atlantic, E: Bay of Biscay & Iberian Coast, F: Canary Islands and G: Barents Sea. Red circle indicates SE hydrophone from the Oceanic Northeast Atlantic. (B) Spectrogram (1024-point FFT, Hann window, 50% overlap) of a fin whale song showing the acoustic parameters analysed in this study.

**Figure 2.**
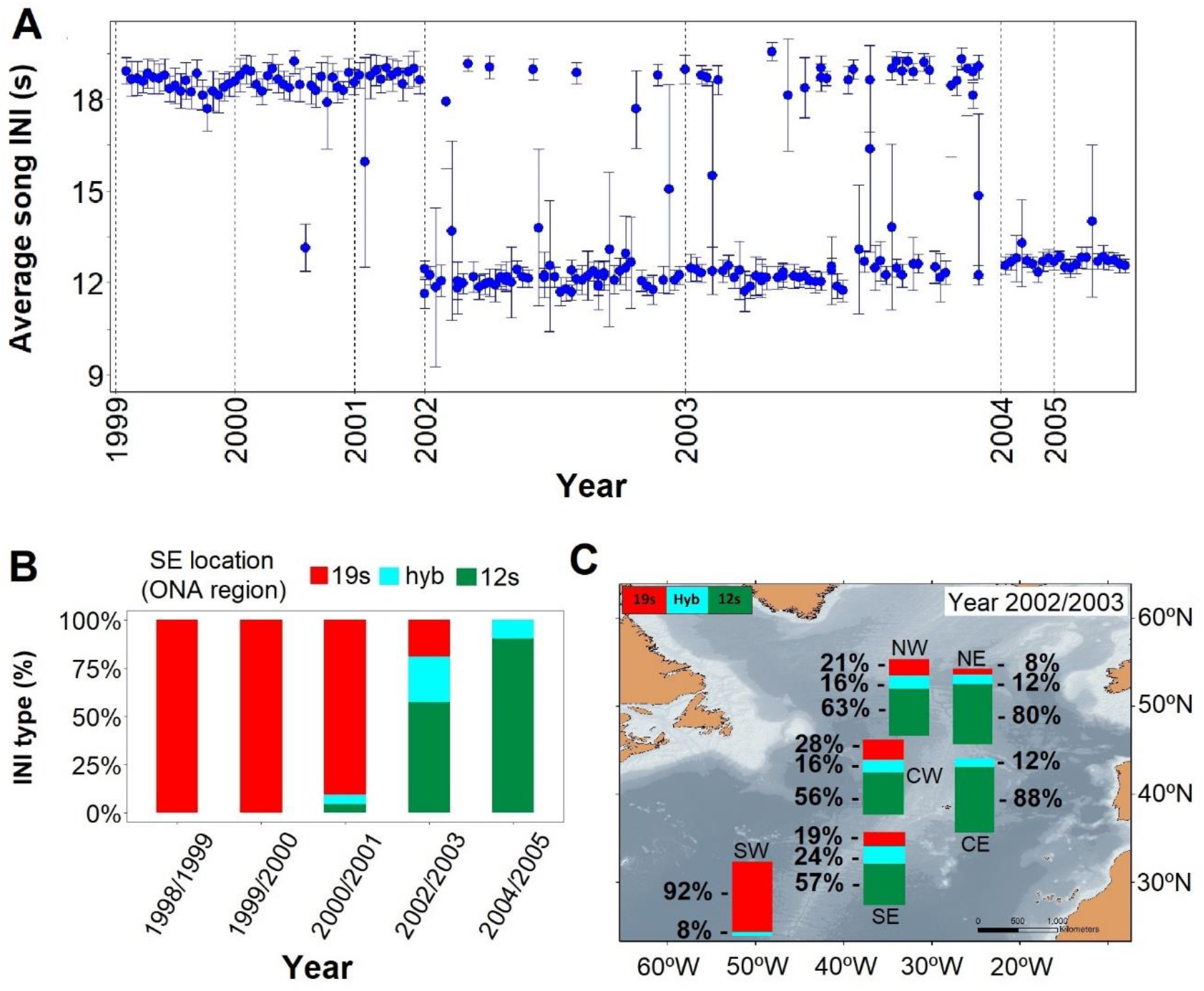
(A) Inter-note intervals (INIs) from 1999 to 2005 for the ONA region (B) Percentage of songs with each INI type (19s, hyb -hybrid, 12s) in the SE hydrophone from the ONA region during the song shift in 1999 – 2005. (C) Map showing the percentage of songs with each INI type for locations within the ONA region hydrophones during the 2002/2003 singing season.

**Figure 3.**
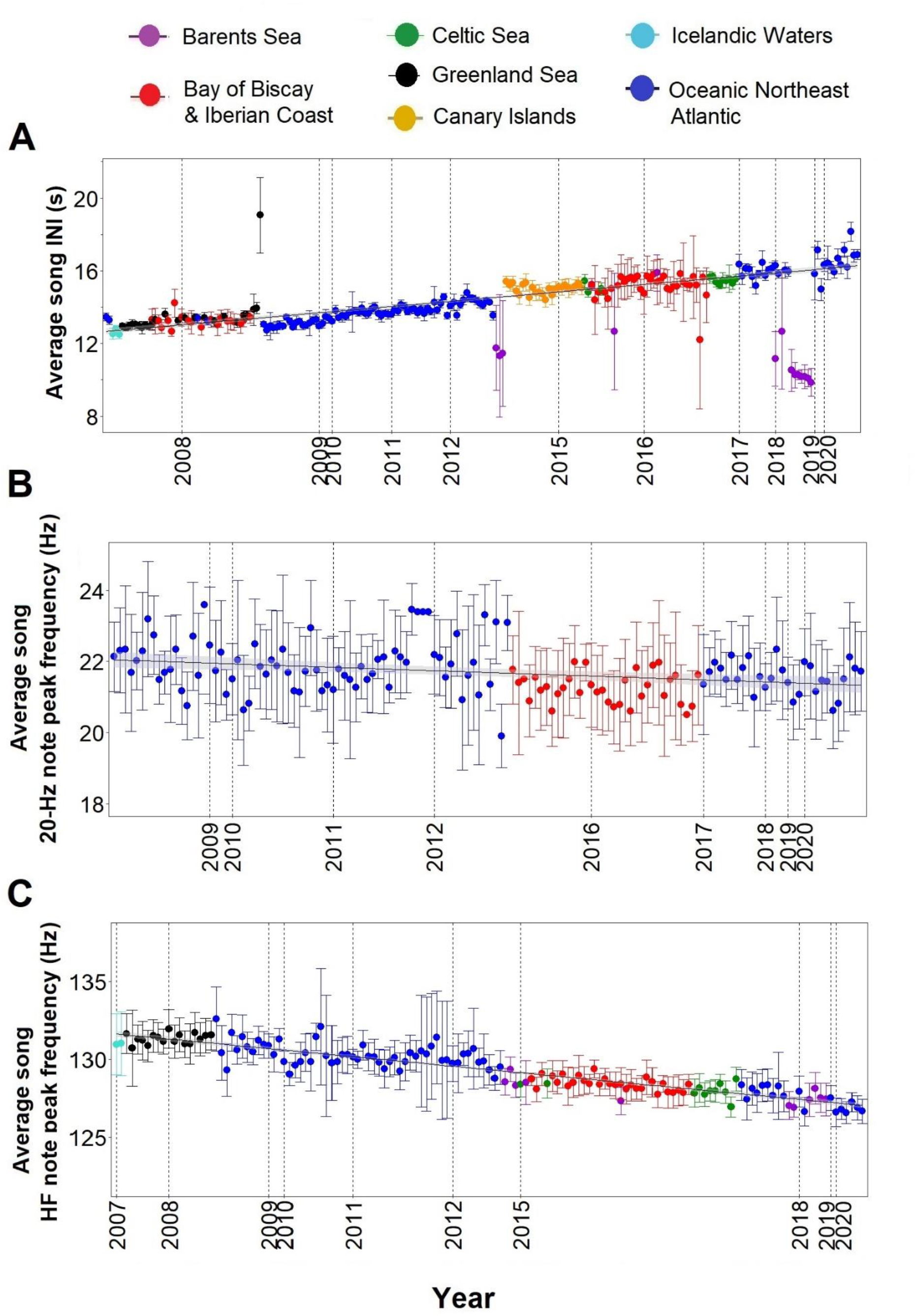
(A) Inter-note intervals (INIs) from 2007 to 2020 for all regions sampled. (B) Peak frequencies of the 20-Hz note for BBIC and ONA regions sampled with Ecologic Acoustic Recorders (2008 – 2020). (C) Peak frequencies of the High Frequency (HF) note for all regions sampled. Points represent average values per song, error bars are standard deviations and black lines represent the fitted linear regression model with confidence intervals in shadowed grey.

### Gradual song changes and conformity

After the song transition, we found a gradual change in three fin whale song parameters over a period of 13 years with all regions fitting the trend. The only exception was the Barents Sea and some songs from the Bay of Biscay & Iberian Coast (BIIC) region in 2016/2017 where INIs differed from the rest of the sampled area. From 2007 to 2021, INIs increased at 0.26 s/yr (Adj. R-sq.= 0.7; *p*<0.001) (Fig. 3A). For this note, peak frequencies decreased at an almost negligible rate of −0.06 Hz/yr (Adj. R-sq.= 0.1; *p*<0.001) (Fig. 3B) while peak frequencies of the HF note decreased at a rate of −0.35 Hz/yr (Adj. R-sq.= 0.8; *p*<0.001) with all regions fitting the trend, including the Barents Sea (Fig. 3C).

When comparing data from different regions (Icelandic Waters, Greenland Sea, ONA, BBIC, Canary Islands, Barents Sea and Celtic Sea) with simultaneous recordings (i.e., in the same singing season) results showed unimodal overlapping distributions in INIs and HF note peak frequencies. The only exception was the Barents Sea region, where INIs differed from the Canary Islands in 2014/2015 (Barents Sea: ~ 9s and ~ 14s; Canary Islands: ~ 15s) and from the ONA region in 2017/2018 (Barents Sea: ~ 10s; ONA: ~16s) (Fig. 4). In 2015/2016, the BIIC region had 21% of hybrid songs that included a small number of 9s INIs (identified only in songs from the Barents Sea region) but showed no hybrid songs back in 2007/2008 (Fig. 3A and Fig. 4A).

**Figure 4.**
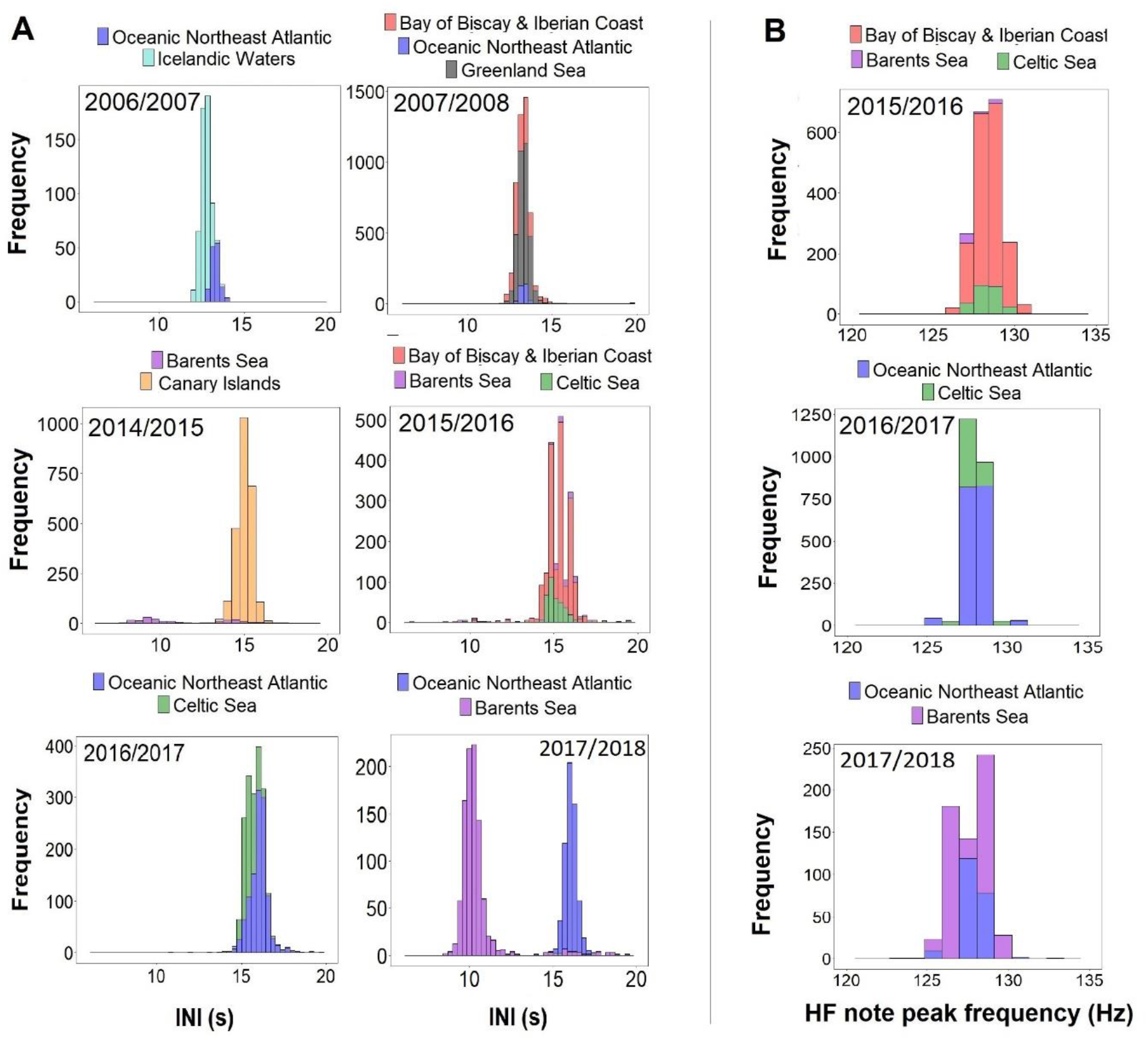
Histograms of inter-note intervals (INIs) and higher frequency (HF) note peak frequencies by singing season (Oct-Mar) from regions with concurrent data.

## DISCUSSION

The changing pattern of different fin whale song parameters reported here for a wide area of the Central and Eastern North Atlantic provides evidence of vocal learning and conformity in this species. Moreover, decoupled variations in rhythm and frequency (i.e. INIs change rapidly but frequencies do not) and between note types (negligible decrease in frequencies of 20-Hz note compared to marked decrease of HF note) reveal the complex interplay between different selective pressures and suggest distinct functions for these song parameters and components.

The dynamics of rapid replacement of fin whales’ song types described here for the ONA region cannot be explained by an environmental causation. The shift in INIs found in the ONA region seemed to occur simultaneously at northern feeding grounds, in the so-called Northeast North Atlantic (NENA) region^22^. This variation in INI patterns during the same singing season between neighbouring locations within ONA, together with an identical shift in INIs documented for the same period in the environmentally distant NENA region^22^, strongly suggest that the transition in INI variants was not a response to local acoustic environments. A population replacement as a potential explanation for the shift in song types also seems unlikely. If the change in INIs (from 19s to 12s) resulted from the replacement of one population by another, we would not find hybrid songs containing both INIs during the transition period, as the new song pattern would simply substitute the former, as documented for fin whale songs off Southern California^25^. Instead, we suggest that the rapid and complete turnover of fin whale song types along a spatial gradient, with all males adopting the new song (i.e., conformity), and the existence of hybrid songs, is the result of cultural transmission, the social learning of information or behaviours from conspecifics^37^.

The geographic variation in bird songs is primarily attributed to their ability of learning to vocalize through imitation^40^. Fin whale song INIs are also regionally distictive^22^ and all males within a certain area conform to the same INI ^24,25,27^ (*this study*). In addition, differences in fin whale songs among regions do not reflect estimates of genetic divergence^22^, further suggesting that song rhythm may be socially learned in this species. Learning of novel rhythms (i.e., INIs) is considered vocal usage learning because existing signals are given in a new sequence^4^, but has not yet been demonstrated in animals despite evidence of novel rhythm imitation^16^. For example, sperm whales (*Physeter macrocephalus*) are able to match their clicks to the rhythm of a ship depth sounder^38^ and use codas (i.e., rhythmic patterns of clicks) for communication that are unique to each vocal clan^39^ and may be socially learned^39^. Moreover, fin whales can also sing songs from other populations that differ in note composition^26^, providing support for vocal production learning in this species. Co-occurrence of both vocal learning strategies (usage and production learning) are not rare and has been documented in several species^16^.

Fin whale song INIs not only varied rapidly but also gradually. After the song transition, we found a gradual increase in song INIs along with a decrease in peak frequencies of the 20-Hz and HF notes. These findings are in line with the gradual trends of decreasing frequencies^27,29,41^ and increasing INIs^25,27,30^ described for fin whale songs in other ocean basins and in the Mediterranean Sea. Contrarily to the rapid changes in INIs, a global-scale process of cultural transmission cannot explain these directional changes. First, changes in INIs and frequencies occur at different rates in different oceans and there is no convergence in song acoustic characteristics across populations ^25,27,28^. And second, a similar pattern of decreasing frequencies and increasing INIs has been described for blue whale (*Balaenoptera musculus*) songs^42,43^, and decreasing frequencies have been reported for bowhead whales (*Balaena mysticetus*) calls^44^. Such gradual song changes in multiple species and different ocean basins suggest an adaptation to a common selective pressure, which does not mean that within-region conformity in song characteristics does not arise from cultural transmission (see below). So far, none of the proposed hypotheses can convincingly explain the slow evolution in fin whale songs^42,44^ partly because they lack a global approach. Large-scale and long-term datasets would help understanding if fin whale song INIs and frequencies are constantly evolving or started changing recently in response to a new driver.

Our results also show that variations in INIs and frequencies are uncoupled because i) the rapid shift in INIs was not matched by a similar shift in 20-Hz note frequencies and ii) frequencies of the HF note frequency decreased six times faster than the 20-Hz note. These findings suggest distinct functions and selective pressures acting on different acoustic parameters and song components. Fin whale song INIs are geographically distinct^22^ and have been used to differentiate stocks and populations^23–25^. Assuming the rapid song change reported here is the result of cultural transmission, fin whales singing the two song types (12s and 19s) would need to be in acoustic contact for the song transfer to occur. If so, the mixing of these two populations have resulted in the rapid adoption of just one single song type. This rapid replacement of song types resembles humpback whale song revolutions, in which a population’s song is rapidly replaced by a novel song type introduced from a neighbouring population^12^. Yet, more data would be needed to prove the novelty and origin of the 12s fin whale song to unequivocally confirm a cultural revolution in fin whale songs. In any case, both song plasticity (i.e., learning the new song) and conformity seem to be selected in humpback whale song evolutions^13^ and it is possible the same occurs for fin whale song INIs. Conversely, fin whale song frequencies have a limited variation compared to INIs. Fundamental frequencies of this specie’s songs seem especially adapted for long-range communication in pelagic environments. Fin whales disperse across deep open waters during their breeding season^45,46^ and their song frequencies match a particular frequency band with low levels of noise in deep waters (i.e., quiet window)^47,48^. Humpback and right whales (*Eubalaena* sp.) aggregate in coastal breeding grounds ^49,50^ and use higher frequency songs and calls that better transmit in shallow environments (quiet window: 100-400 Hz)^51^ and do not need to reach distant conspecifics^48,52^. Therefore, the acoustic environment during the mating season could constrain variation in song frequencies to keep them within the quiet window. Yet, these frequency variations are far more limited for the 20-Hz note, that show a negligible changing rate, than for the HF note. Differing changing trends between song components has also been found for two song units of the Sri Lankan pygmy blue whale (*Balaenoptera musculus brevicauda*) that authors attributed to distinct functions, arguing that the lower frequency component is more conserved because it conveys information on species identity^53^. In some bird species, specific segments of their songs remain relatively constant over decades, possibly reflecting their role in defining the species^54,55^. In the same way, the fin whale 20-Hz note produced worldwide^17,19,56–58^ may encode species identity, while the HF note, which varies geographically at a larger scale than INIs and is not used by all populations^22,58^, may be more prone to variation. This is further supported by our results showing that fin whales from the distant Barents Sea region differ from the rest of the sampled area in their song INIs but not in their HF note frequencies, that fit within the general trend. Therefore, fin whale song INIs are indicative of acoustic populations while frequencies may respond to changes in the acoustic environment.

The adoption of rapid and gradual changes in three song parameters by all males across a wide area of the Central and Eastern North Atlantic indicates strong song conformity in fin whales. Song conformity seems to be common in fin whales worldwide^24–28,30,59^, which suggests it may be relevant for the function of their songs. Male fin whale songs are believed to act as mating displays because all biopsied singing whales were males^18^ and singing intensifies during the specie’s breeding season^17,19–21^. In bird species that use songs as acoustic displays, song conformity may be driven by a female preference for the most common variant. Females may prefer the local song to a foreign one because it indicates the male’s copying abilities or male’s genes better adapted to the local environment^60^. In fin whales, conformity in song INIs and frequencies could provide the same advantages and even assist females assessing male quality. The fact that fin whale singing decreases when they swim faster has led authors to hypothesize that singing while swimming may be an indicator of a male’s stamina^61^. If all males sing the same song, it may be easier for females to assess differences between individual male’s song characteristics.

The low-frequency, high amplitude, repetitive and simple characteristics of fin whale songs seem especially suited for long-range communication ^48,62^, which may be an adaptation to the dispersed and pelagic distribution of this species during the breeding season^45,46^. Hence, songs may also be used for ranging of conspecifics and perhaps, environmental sensing. Evidence suggests that baleen whales are capable of both. Modelling indicates that songs from humpback whales could be used as long-range sonar^63^. Bowhead whales showed echo-ranging behaviour when navigating under heavy ice conditions during their spring migration^64^. Blue whales changed their usual calls in association with sudden changes in oceanographic conditions^65^. Finally, acoustically tracked fin and Bryde’s whales (*Balaenoptera edeni*) clearly kept parallel tracks suggesting these species are able of ranging conspecifics^66,67^

Estimating the distance to a source involves assessing signal degradation through frequency-dependent attenuation and environmental echoes (i.e., reverberation)^68^. This process requires knowledge of both the environmental conditions and the undistorted source signals^69^. Additionally, an animal needs to be familiar with the time-varying features of a specific sound to be able to accurately judge the distance^70^. This way, we can hypothesize that conformity in fin whale song INIs and frequencies may be partially associated with the necessity of this species to sense their environment and conspecifics through time-frequency dispersions of their own and other whale’s songs. In fact, frequency conformity in blue whale songs have been hypothesized to facilitate ranging abilities by using the Doppler effect^71^. Also, Bryde’s whales synchronised their calls when in close distance and kept parallel tracks while the synchronisation lasted^67^. As with songbirds, vocal learning in cetaceans may have evolved, partly, because it enabled ranging sound sources more successfully^70,72^.

Results from this and other studies suggest that male fin whales are in acoustic contact over vast areas and adjust their song properties to match those of conspecifics^27,28,59^. These acoustic communities culturally evolve more quickly and efficiently than genetic communities^22^ and should be considered in conservation strategies when delimiting stocks or populations. These results have also implications for cue counting approaches, that use cue rates to convert density of sounds to density of animals^73^. The temporal and spatial changes in fin whale song INIs found here affect cue rates and need to be accounted for to avoid bias in estimating densities using passive acoustic monitoring in this species. Finally, understanding the cultural evolution of fin whale songs can inform on the species’ ability to adapt, or not, to a changing environment. The unique large spatial scale over which fin whales communicate, although technologically challenging for researchers, opens interesting perspectives in the processes of animal acoustic communication.

## Materials and methods

### Sampling locations

Acoustic data were collected from 15 locations in the Central and Northeast Atlantic Ocean, grouped into seven regions as defined by the International Council for the Exploration of the Sea (ICES)^31^: Greenland Sea, Icelandic Waters, Celtic Sea, ONA, BBIC, Canary Islands and Barents Sea (Supplementary Table S1, Fig. 1A).

### Data collection

Recordings from 1999 to 2020 collected by different research groups with varied objectives were compiled and standardised. Not all regions were sampled in all years and time periods. Recordings were either continuous or duty-cycled, with sampling rates ranging from 100 Hz to 48 kHz (Supplementary Table S1 and Fig. S1). Ocean-Bottom Seismometers (OBS) were used in the Canary Islands (2014-2015) and in the BBIC (2007-2008). The OBS channel with the highest signal-to-noise ratio (SNR) was used in the analysis. The hydrophone channel was selected for recordings in the Canary Islands, while the seismometer channel (vertical component Z) was preferred for recordings from the BBIC (2007-2008). Fixed autonomous recorders (AR) were used in the remaining regions (Supplementary Table S1).

### Song selection criteria

We focused the analyses on data collected between October and March (hereafter singing season), because fin whale song parameters show less variation during this period^22^ and seasonal variation was outside the scope of this study. All datasets were manually inspected to identify days with fin whale 20-Hz notes^17^ or double pulsed songs containing the 20-Hz and higher frequency note (hereafter HF note)^22^ (Fig. 1B). For the Azores dataset, a Low Frequency Detection and Classification System (LFDCS)^32^ was used following procedures described in Romagosa et al. (2020)^33^. Spectrograms of days with fin whale detections were manually analysed using Adobe Audition 3.0 software (Adobe Systems Incorporated, CA, USA) to select periods with good quality notes, based on: a) a high SNR song, b) absence of masking from noise, c) presence of a single singer and d) occurrence of notes organized in long series. The last criterion could not be applied in recordings with small duty cycles (SW Portugal 2015-2016, Azores 2008-2011 and North and South Porcupine) (Supplementary Table S1); nevertheless, regularly spaced notes could still be identified as part of songs.

### Song sampling

Selected days with detections were non-consecutive to minimize sampling the same animal multiple times. The number of sampled days varied depending on the quality of fin whale songs found in the recordings. The average number of days sampled per singing season was 10.8 days, and the average number of notes analysed per song was 108 (Supplementary Fig. S1). Recordings from the Canary Islands, BBIC (2007-2008), and ONA regions, except for the Azores (Supplementary Table S1), were excluded from the analysis of the HF note, because sampling rates were too low to enable detection of the HF note frequency (~130-Hz)^22^ (Supplementary Fig. S1).

### Measurement of song parameters: inter-note intervals and peak frequencies

Selected days with good quality notes were fed into a band-limited energy detector in Raven Pro 1.5 software (Cornell Lab of Ornithology, Ithaca, NY, USA) that automatically selected all 20-Hz and HF notes in the spectrogram. All selections were checked manually by the same analyst to ensure that notes were well imbedded in the selection square. Spectrogram characteristics were adjusted to visualise all data with the same frequency and time resolution. For each selected note, the software measured begin and end time, and peak frequency. Inter-note intervals (INIs) were calculated by subtracting the time difference between the begin time of a 20-Hz note and the begin time of the following 20-Hz note^17,25^ (Fig. 1B). Peak frequencies were measured for 20-Hz and HF notes and represent the value at which the maximum energy in the signal occurs. It is considered a robust measurement because it is based on the energy within the selection and not the time and frequency boundaries of the selection^34^. Only one sequence of notes or song fragment (hereafter referred as song) was analysed per day in each location. If multiple songs were found in one day, the one with the highest SNR was selected. For each song, we calculated the mean and standard deviation of INIs and of peak frequencies of the 20-Hz and HF notes.

### Transition between song INIs

Temporal and spatial changes in song INIs were investigated by using data from the ONA region, the only area with recordings during the song transition period (2000-2005). The SE ONA hydrophone was used to investigate how song types changed over time (Fig. 1A; red circle), by calculating the percent number of each song type identified in each singing season (October-March) from 1998/1999 to 2004/2005. Spatial variation in song types was examined by comparing the percentage of different song types from six ONA locations (NE, NW, CE, CW, SE and SW) that covered part of the transition period (2002/2003) (Fig. 1A and Supplementary Fig. S1).

### Trends in song properties

Data from all regions were plotted in chronological order to investigate how song parameters varied over time. A linear regression model was fit to each response variable (INIs and peak frequencies of the 20-Hz and HF notes) using a Gaussian distribution and year as the explanatory variable. Model assumptions were verified by plotting residuals versus fitted values and residual QQ plots to check for homogeneity of variance and normality (Supplementary Figs. S2A, B & C). In the case of INIs, the model was fitted using data from all regions between 2007 and 2020, except from the Barents Sea, that showed different INIs. (Fig. 3A). Measurements of 20-Hz peak frequencies were greatly affected by the recording equipment (Suppl. Information and Fig. S3). For this reason, only data from the Ecological Acoustic Recorders (EARs)^35^, which sampled the longest period (2008 – 2020) (Supplementary Fig. S1), were used to explore temporal variations in the peak frequencies of the 20-Hz note (Fig. 3B). All statistical analyses were performed in R (v. 4.0.2)^36^.

### Regional comparison of song parameters

Given inter-annual variations in fin whale song parameters^25,27^, only songs recorded within the same singing season were used to compare song parameters among regions. Histograms were built for each singing season to investigate differences in the distribution of INIs and peak frequencies of the HF note per region sampled.

## ACKOWLEDGEMENTS

Research was supported by the Portuguese Science & Technology Foundation (FCT), the Azorean Science & Technology Fund (FRCT) and the EC through research projects TRACE-PTDC/MAR/74071/2006, MAPCET-M2.1.2/F/012/2011, FCT-Exploratory-IF/00943/2013/CP1199/CT0001, AWARENESS-PTDC/BIA-BMA/30514/2017, co-funded by FEDER, COMPETE, QREN, POPH, ESF, Lisbon and Azores Regional Operational Programme, Portuguese Ministry for Science and Education. Okeanos R&D Centre is supported by FCT through the strategic fund (UIDB/05634/2020). Canary Island data was provided by the Institut de Ciències del Mar under the ‘Severo Ochoa Centre of Excellence’ accreditation (CEX2019-000928-S). Data used from the ONA region is a NOAA-PMEL contribution number 5326. The Celtic Sea data belongs to the ObSERVE Acoustic project, initiated and funded by the Department of Communications, Climate Action and Environment in partnership with the Department of Culture, Heritage and the Gaeltacht under Ireland’s ObSERVE Programme. Data collection in the Barents Sea region was made by the Italian CNR under the Arctic Field Grant project KUAM (235878/E10), funded by the Norwegian Research Council through the Svalbard Science Forum, and under the Project Calving SEIS (244196/ E10) funded by the Norwegian Research Council. Vesterålen data was provided by the LoVe Ocean Observatory project, led by the Institute of Marine Research and funded by the Norwegian Research Council and Equinor. Iceland data was collected under Velux Fonden and Knud Højgårds Fond funding. M.R. was supported by a DRCT doctoral grant (M3.1.a/F/028/2015). IC was supported by the FCT-IP Project UIDP/05634/2020. A.P. was supported by project AWARENESS “PTDC/BIABMA/30514/2017” and “UIDB/50019/2020–IDL”. T.A.M. by CEAUL (funded by FCT – Fundação para a Ciência e a Tecnologia, Portugal, through the project UIDB/00006/2020) and the LMR ACCURATE project (contract no. N3943019C2176). R.P. was supported by an FCT grant (SFRH/BPD/108007/2015). M.A.S. was funded by FCT (IF/00943/2013), EC (SUMMER H2020-EU.3.2.3.1, GA 817806) and the Operational Program AZORES 2020, through the Fund 01-0145-FEDER-000140 “MarAZ Researchers: Consolidate a body of researchers in Marine Sciences in the Azores” of the European Union. We are grateful to Marc Lammers, for providing the EARs and technical support, and to Sérgio Gomes, Norberto Serpa and all skilled skippers and crew that participated in the preparation and deployment of the EARs at DOP/IMAR and all other instruments used in this study.

## AUTHOR CONTRIBUTIONS

Conceptualization, M.R., S.N. and M.A.S.; Methodology, M.R., S.N., T.A.M. and M.A.S.; Data analysis, M.R.; Data collection, M.R., S.N., I.C., R.D., J.R., J.O., D.M., A.P., A.U., E.P., S.A., G.B., L.M., M.RA, R.P. and M.A.S.; Funding acquisition: M.R., R.P. and M.A.S.; Writing – Original Draft, M.R. and M.A.S.; Writing – Review & Editing, M.R, S.N, I.C, T.A.M, R.D, J.O, D.M, A.P, A.U, E.P, S.A., M. R, L.M, M.RA, R.P and M.A.S.

## DATA AVAIALBILITY

Datasets and R scripts used in this study are available from the Dryad Digital Repository: https://doi.org/10.5061/dryad.f1vhhmh13 [74].

