## Supplementary information, tables and figures for "Fin whale singalong: evidence of song conformity"

### SUPPLEMENTARY MATERIAL

*Table S1. Position, recording equipment, depth, duty cycle and sampling rate for each location sampled.*

| Regions | Location | Latitude and longitude (°) | Recording equipment (AR or OBS) | Aprox. depth (m) | Duty cycle on/off (min) | Sampling rate (Hz) |
|---|---|---|---|---|---|---|
| **Greenland Sea** | SE Greenland | 60° N<br>35° W | AR[74] | 800 | Continuous recording | 2000 |
| **Icelandic Waters** | SE Iceland | 64.185° N<br>14.686° W | AR[75] | 60 | Continuous recording | 4000 |
| **Celtic Sea** | North Porcupine | 52.6221° N<br>15.3045° W | AR<br>(AMARs, JASCO Applied Sciences, Halifax, Canada) | 1700 | 2/18 | 32000 |
| | South Porcupine | 49.5478°N<br>13.3723°W | | | | |
| **Oceanic Northeast Atlantic** | North East (NE ONA) | 49.8554°N<br>25.4541°W | AR[74] | 4200 | Continuous recording | 260 |
| | North West (NW ONA) | 47.5941°N<br>-32.4500°W | | 4100 | Continuous recording | |
| | Central East (CE ONA) | 40.3365°N<br>25.0350°W | | 3300 | Continuous recording | |
| | Central West (CW ONA) | 42.7188°N<br>34.7226°W | | 3970 | Continuous recording | |
| | Azores | 38.5396°N<br>29.0434°W | AR (EARs)[72] | 200-400 | 60/138<br>60/210<br>360/1440 | 2000 |
| | South East (SE ONA) | 32° N<br>35° W | AR[74] | 926 | Continuous recording | 110 |
| | South West (SW ONA) | 26°N<br>50°W | | | | |
| **Bay of Biscay & Iberian Coast** | SW Portugal | 35.7798°N<br>10.3584°W | OBS (Silva, S. 2017) | 1993-5100 | Continuous recording | 100 |
| | | 36.5753° N<br>11.5969° W | AR (EARs)[72] | 255 | 3/12 | 2000 |

| | | | | | | |
|---|---|---|---|---|---|---|
| **Canary Islands** | Lanzarote | 28.8997°N 13.2003°W | OBS[76] | 1350 | Continuous recording | 100 |
| **Barents Sea** | West Svalbard | 79.0536°N 11.5471°E | AR (SM2, Wildlife Acoustics, US) | 75 | Continuous recording | 48000 |
| | Vesterålen, Norway | 68.9°N 14.3838°E | AR (SB35 ETH, Ocean Sonics) | 258 | Continuous recording | 32000 |

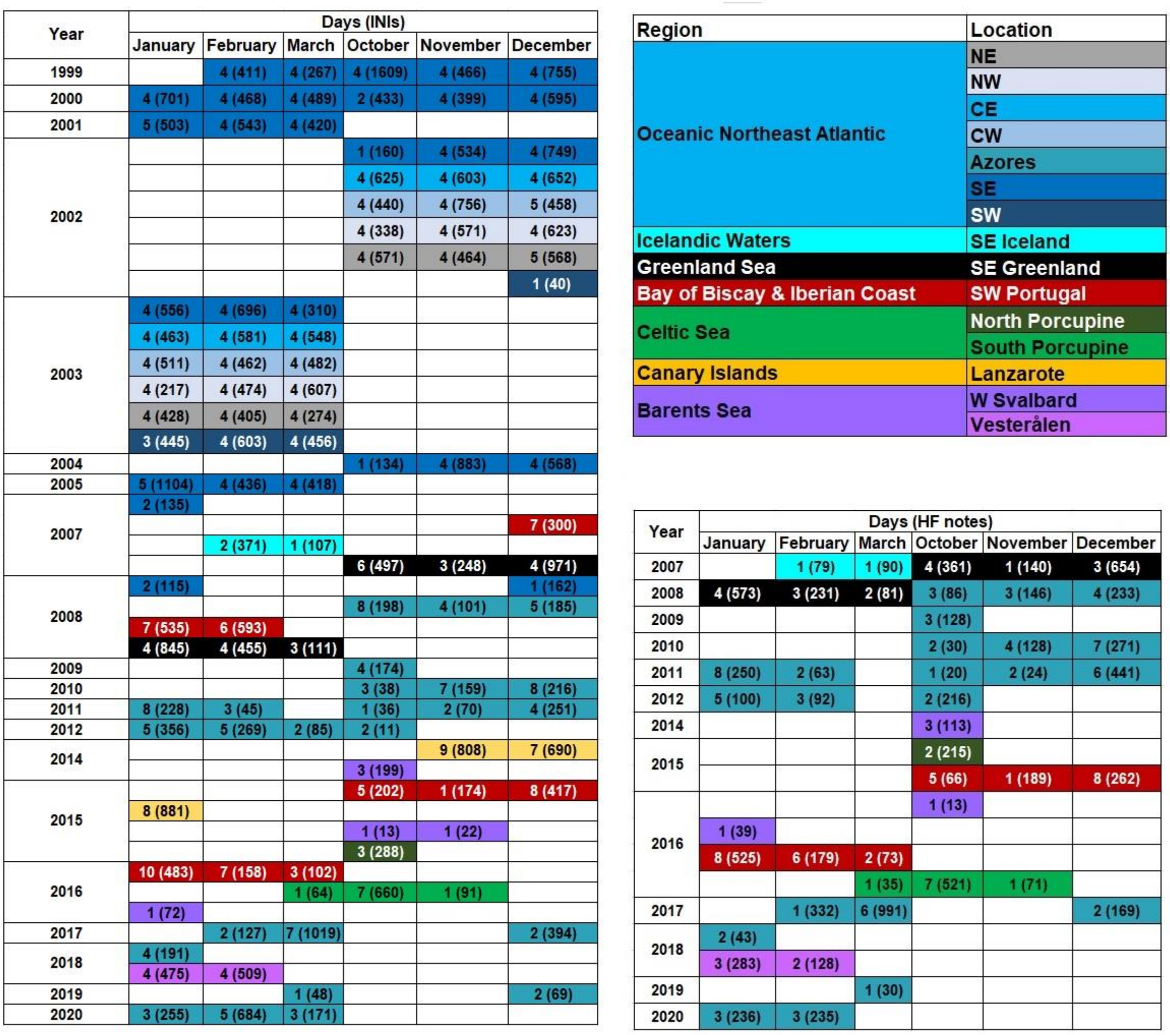

| Year | Days (INIs) | | | | | |
|---|---|---|---|---|---|---|
| | January | February | March | October | November | December |
| 1999 | | 4 (411) | 4 (267) | 4 (1609) | 4 (466) | 4 (755) |
| 2000 | 4 (701) | 4 (468) | 4 (489) | 2 (433) | 4 (399) | 4 (595) |
| 2001 | 5 (503) | 4 (543) | 4 (420) | | | |
| 2002 | | | | 1 (160) | 4 (534) | 4 (749) |
| | | | | 4 (625) | 4 (603) | 4 (652) |
| | | | | 4 (440) | 4 (756) | 5 (458) |
| | | | | 4 (338) | 4 (571) | 4 (623) |
| | | | | 4 (571) | 4 (464) | 5 (568) |
| | | | | | | 1 (40) |
| 2003 | 4 (556) | 4 (696) | 4 (310) | | | |
| | 4 (463) | 4 (581) | 4 (548) | | | |
| | 4 (511) | 4 (462) | 4 (482) | | | |
| | 4 (217) | 4 (474) | 4 (607) | | | |
| | 4 (428) | 4 (405) | 4 (274) | | | |
| | 3 (445) | 4 (603) | 4 (456) | | | |
| 2004 | | | | 1 (134) | 4 (883) | 4 (568) |
| 2005 | 5 (1104) | 4 (436) | 4 (418) | | | |
| 2007 | 2 (135) | | | | | |
| | | | | | | 7 (300) |
| | | 2 (371) | 1 (107) | | | |
| | | | | 6 (497) | 3 (248) | 4 (971) |
| 2008 | 2 (115) | | | | | 1 (162) |
| | | | | 8 (198) | 4 (101) | 5 (185) |
| | 7 (535) | 6 (593) | | | | |
| | 4 (845) | 4 (455) | 3 (111) | | | |
| 2009 | | | | 4 (174) | | |
| 2010 | | | | 3 (38) | 7 (159) | 8 (216) |
| 2011 | 8 (228) | 3 (45) | | 1 (36) | 2 (70) | 4 (251) |
| 2012 | 5 (356) | 5 (269) | 2 (85) | 2 (11) | | |
| 2014 | | | | | 9 (808) | 7 (690) |
| | | | | 3 (199) | | |
| 2015 | | | | 5 (202) | 1 (174) | 8 (417) |
| | 8 (881) | | | | | |
| | | | | 1 (13) | 1 (22) | |
| | | | | 3 (288) | | |
| 2016 | 10 (483) | 7 (158) | 3 (102) | | | |
| | | | 1 (64) | 7 (660) | 1 (91) | |
| | 1 (72) | | | | | |
| 2017 | | 2 (127) | 7 (1019) | | | 2 (394) |
| 2018 | 4 (191) | | | | | |
| | 4 (475) | 4 (509) | | | | |
| 2019 | | | 1 (48) | | | 2 (69) |
| 2020 | 3 (255) | 5 (684) | 3 (171) | | | |

| Region | Location |
|---|---|
| Oceanic Northeast Atlantic | NE |
| | NW |
| | CE |
| | CW |
| | Azores |
| | SE |
| | SW |
| Icelandic Waters | SE Iceland |
| Greenland Sea | SE Greenland |
| Bay of Biscay & Iberian Coast | SW Portugal |
| Celtic Sea | North Porcupine |
| | South Porcupine |
| Canary Islands | Lanzarote |
| Barents Sea | W Svalbard |
| | Vesterålen |

| Year | Days (HF notes) | | | | | |
|---|---|---|---|---|---|---|
| | January | February | March | October | November | December |
| 2007 | | 1 (79) | 1 (90) | 4 (361) | 1 (140) | 3 (654) |
| 2008 | 4 (573) | 3 (231) | 2 (81) | 3 (86) | 3 (146) | 4 (233) |
| 2009 | | | | 3 (128) | | |
| 2010 | | | | 2 (30) | 4 (128) | 7 (271) |
| 2011 | 8 (250) | 2 (63) | | 1 (20) | 2 (24) | 6 (441) |
| 2012 | 5 (100) | 3 (92) | | 2 (216) | | |
| 2014 | | | | 3 (113) | | |
| 2015 | | | | 2 (215) | | |
| | | | | 5 (66) | 1 (189) | 8 (262) |
| 2016 | | | | 1 (13) | | |
| | 1 (39) | | | | | |
| | 8 (525) | 6 (179) | 2 (73) | | | |
| | | | 1 (35) | 7 (521) | 1 (71) | |
| 2017 | | 1 (332) | 6 (991) | | | 2 (169) |
| 2018 | 2 (43) | | | | | |
| | 3 (283) | 2 (128) | | | | |
| 2019 | | | 1 (30) | | | |
| 2020 | 3 (236) | 3 (235) | | | | |

*Fig. S1. Number of non-consecutive days sampled and INIs measured (in brackets) (left), and number of non-consecutive days sampled and HF notes measured (in brackets) (right), for each location and year.*

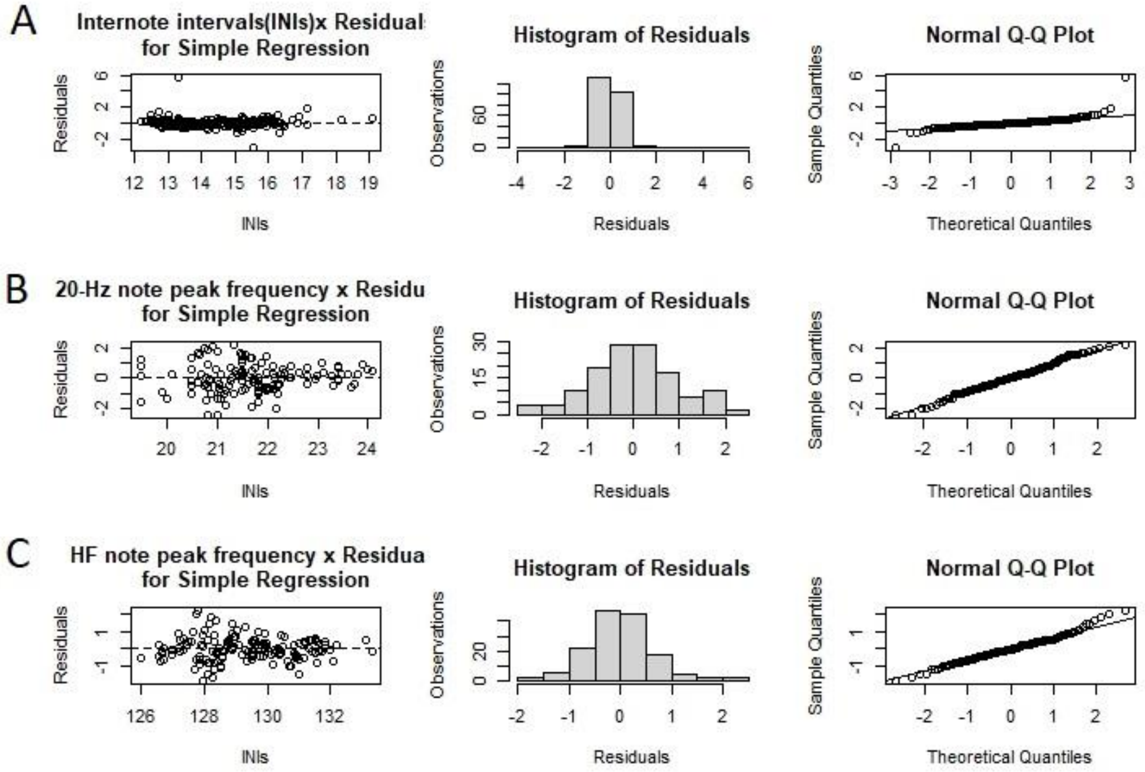


Figure S2. Linear model validation plots for INIs (A), 20-Hz note peak frequencies (B) and HF note peak frequencies (C).

#### *1. The effect of recording equipment on fin whale song parameters*

When comparing data from multiple sensors an obvious question is whether the results might be dependent on the specific sensors considered. To investigate the influence of the acoustic recorder type, OBS or ARs, on the song parameters, we analysed the same song fragment, consisting of 209 notes, recorded by the hydrophone channel of an OBS and an AR, specifically an Ecological Acoustic Recorder (EAR)[72]. The two instruments were deployed at ~6 km from each other in the Azores region in spring of 2019. Measurements of INIs and 20-Hz peak frequencies of songs recorded by each instrument were compared using a non-parametric paired samples Wilcoxon Test. Differences in HF note peak frequencies could not be tested because of limitations in the sampling rate of the OBS. Results showed that median INIs measured from OBS (16.52) and EARs (16.42) were not significantly different (p-value = 0.43) but median peak frequencies of the 20-Hz note (median OBS = 23.4; EARs = 21.1) were (p-value < 0.001). Thus, the use of different recorders did not affect INI measurements but influenced measurements of 20-Hz peak frequencies. The effect of the distance to the source in the analysed fin whale song parameters is included in this study, given than these two recorders were positioned at different distances to the singer. Peak frequencies of the 20-Hz note showed a great variability between equipment types (Fig. S3), which hindered the identification of soft trends (i.e., low changing rate). For this reason, only data from the EARs, the longest dataset (2008 – 2020), were used to study temporal variations of the peak frequencies of the 20-Hz note. All statistical analyses were performed using the software R (v. 4.0.2)[73].

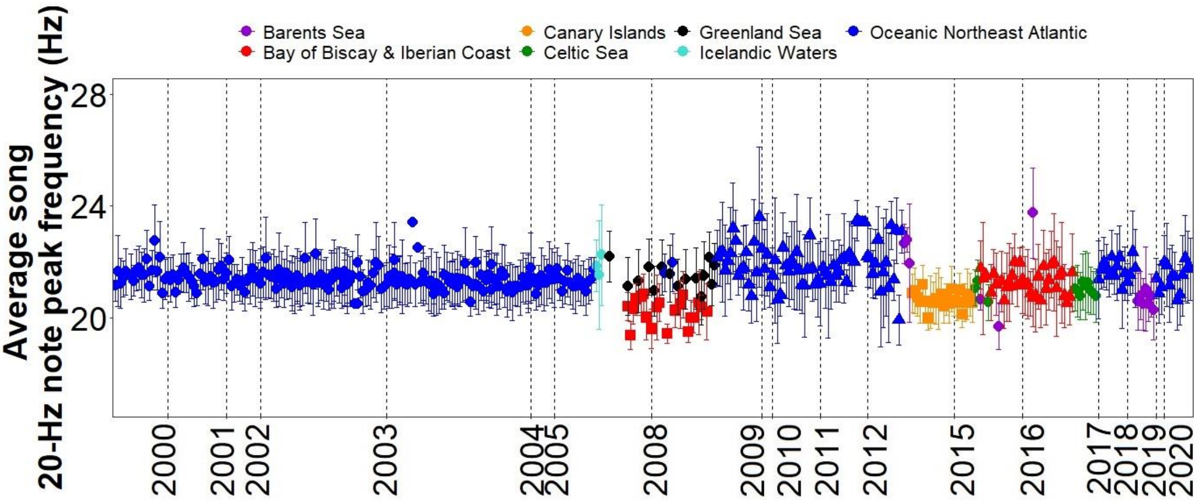


*Figure S3. Peak frequencies of the 20-Hz note for the regions sampled and equipment used: ARs (circle), ARs – EARs (triangle) and OBS (square). Points represent averaged peak frequencies per song and error bars are standard deviations.*
